# Neural Substrates of Duration Serial Dependence: Putamen Suppression of the Prior Duration Trace

**DOI:** 10.64898/2026.05.12.724564

**Authors:** Si Cheng, Siyi Chen, Jiao Wu, Chunyu Qu, Zhuanghua Shi

## Abstract

Perceptual judgments are biased toward recent experience, a phenomenon called serial dependence. For orientation and motion, this carryover has a demonstrated locus in the early visual cortex; for time, the circuits that hold the previous duration are unknown. Using fMRI, we asked where the preceding duration is represented, and whether the same neural mechanism carries recent information about a non-temporal feature of the same stimulus. Participants viewed a moving random-dot display and reproduced either its duration or its direction, retro-cued as to which feature to report, thereby separating the memory trace from the demands of task switching. Duration reproduction was attracted toward the preceding duration, four times more strongly when the same task was repeated. Direction judgments exhibited a much smaller bias that reversed sign with the preceding task. The preceding duration was negatively associated with retro-cue-locked activity in the bilateral putamen, the region where this effect survived whole-brain correction. The estimate held after accounting for both current and preceding responses. Direction produced no comparable signal anywhere. Switching from the direction task to the time task recruited the inferior frontal junction and dorsolateral prefrontal cortex, neither of which carried a preceding-duration effect. The prior-duration signal, therefore, localized to the dorsal striatum, distinct from the frontoparietal regions in task reconfiguration. These findings argue against a unitary neural mechanism for serial dependence and suggest that recent-history biases are likely domain-specific: for perceived time, the critical trace resides in the striatum rather than in the visual cortex.

## 1. Introduction

Perceptual judgments are seldom made in isolation. Even when observers evaluate each stimulus independently, their responses are systematically biased by recent history, a phenomenon termed *serial dependence* (Fischer and Whitney, 2014). Current estimates are typically attracted toward recently encountered stimuli, producing a smoothing effect across successive episodes. This attractive bias has been documented across orientation (Cicchini et al., 2017; Fischer and Whitney, 2014), numerosity (Cicchini et al., 2014), face identity (Liberman et al., 2014), and time perception (Cheng et al., 2024a; Glasauer and Shi, 2022). Glasauer and Shi (2022) demonstrated that individual differences in sequential bias reflect personal beliefs about temporal continuity, suggesting that serial dependence arises from Bayesian integration of current evidence with prior experience. The breadth of these effects raises two questions that behavioral demonstrations alone cannot answer. First, are serial biases carried by a mechanism common to all stimulus dimensions, like magnitude, or by neural systems specific to the judged feature? Second, when the judged dimension changes from one trial to the next, can the neural trace of the previous stimulus be distinguished from the activity of switching to the new judgment? Both require neural evidence, which remains scarce: most neuroimaging work has focused on orientation, leaving the temporal domain largely unexplored.

For orientation, fMRI studies reveal a two-stage architecture. Schwiedrzik et al. (2014) first identified a frontoparietal network supporting perceptual hysteresis, while adaptation effects remained in early visual cortex. Subsequent work confirmed that V1– V3 carries *repulsive* neural shifts consistent with sensory adaptation, yet the behavioral output is attractive (John-Saaltink et al., 2016; Sheehan and Serences, 2022). Sheehan and Serences (2022) resolved this paradox with a computational model showing that an optimal readout mechanism inverts repulsive encoding into an attractive estimate, while Van Bergen and Jehee (2019) demonstrated that decoded sensory uncertainty predicts the magnitude of bias, consistent with Bayesian weighting. Whether this sensory-adaptation-plus-readout architecture applies to *temporal* serial dependence is untested. The only fMRI study of duration serial dependence (Cheng et al., 2024a) implicated the caudate nucleus and hippocampus, memory-related structures rather than sensory cortex, suggesting a fundamentally different neural implementation.

Behavioral evidence for temporal serial dependence is robust: duration reproduction is systematically attracted toward the previous trial’s duration across sub-second to supra-second intervals (Cheng et al., 2024a, 2024b, 2024c; Glasauer and Shi, 2022). Critically, this bias is modulated by task relevance — serial dependence is substantially stronger on task-repeat than task-switch trials when participants alternate between reproducing duration and another feature (Cheng et al., 2024b; Wu et al., 2026), a pattern also observed in other domains (Bae and Luck, 2020). Behavioral data alone, however, cannot determine why switching reduces the bias: the previous temporal trace may itself be weakened, or it may remain intact but exert less influence after the brain reconfigures for a different task. At the neural level, the striato-thalamo-cortical (STC) circuit — comprising the dorsal striatum, thalamus, and SMA — is the strongest candidate substrate, operating within a wider timing network that extends to the superior temporal and parietal association cortices (Merchant et al., 2013; Wiener et al., 2010). The dorsal striatum encodes elapsed intervals via ramping dynamics (Meck et al., 2008; Yin et al., 2022), so a residual temporal trace should leave its signature in this circuit rather than in early sensory cortex.

Direction of motion provides a natural comparison domain. Whereas duration elicits attractive serial dependence attributed to post-perceptual memory integration, motion direction produces repulsive biases consistent with sensory adaptation (Bae and Luck, 2020; Cheng et al., 2024b). Importantly, direction biases have also been reported to be smaller and less sensitive to the preceding task (Cheng et al., 2024b), unlike the duration repeat-over-switch advantage. Two features of the same stimulus show qualitatively different history effects, and no within-participant design has yet tested whether these effects arise from distinct neural substrates.

The present study addresses this gap using fMRI with a dual-feature, serial-dependence design. On each trial, participants viewed a coherent-motion stimulus preceded and followed by random-motion masks and reproduced either its duration or its direction, as instructed by a retro-cue that followed the trailing mask. This retro-cue design, extending Cheng et al. (2024b) to fMRI, ensures both features are encoded during stimulus presentation while introducing the task-repeat versus task-switch manipulation at the retro-cue phase. Parametric modulators tracked neural carryover of previous-trial duration and direction, while task-repeat versus task-switch contrasts characterized reconfiguration-related activity. If the temporal serial dependence reflects a residual trace with the neural circuitry that represents elapsed time, prior-duration carryover should emerge in the striato-thalamo-cortical network and not in the motion-sensitive visual cortex.

## 2. Method

### 2.1 Participants

Thirty participants were recruited from LMU Munich (Mean age: 26.16; 17 females). All were right-handed, had normal or corrected-to-normal vision, and reported no history of neurological or psychiatric disorders. Participants provided written informed consent and received monetary compensation (15 EUR/h). The sample size was determined a priori based on effect sizes from Cheng (2024; Cohen’s *d* > 0.75); a G\**Power analysis (one-sample* t*-test, d* = 0.75, α = .05, 1 - β = .80) indicated a minimum of 19 participants. Two participants were excluded due to incomplete data acquisition and one due to excessive head motion (mean framewise displacement > 0.5 mm), yielding a final sample of 27 participants.

The study was approved by the Ethics Board of the Department of Psychology at LMU Munich (Approval date: 04.04.2022).

### 2.2 Stimuli and Apparatus

Stimuli were generated using PsychoPy 2023.2.3 and displayed via an MRI-compatible ProPixx DLP LED projector (VPixx Technologies Inc., Canada) at a viewing distance of approximately 110 cm. The visual stimulus was a random dot kinematogram (RDK) presented within a circular aperture (∼10.8° visual angle) containing 200 dots (diameter 0.27°) moving at 7°/s. On each trial, two features varied independently: duration (0.8–1.8 s, grand mean = 1.3 s) and direction (16 discrete values at 22.5° spacing, spanning 0°–360°).

### 2.3 Design and Procedure

The task is illustrated in Figure 1. On each trial, participants observed a coherent motion stimulus and subsequently reproduced either its duration or its motion direction, as indicated by a retro-cue. Each trial began with a fixation cross (0.5 s), followed by a random-motion mask (0.4–0.6 s). The coherent motion stimulus then appeared for its target duration (0.8–1.8 s), followed by a second random-motion mask (0.8–1.6 s). A retro-cue, a letter "T" (time) or "D" (direction), then appeared for 0.5 s, after which participants had up to 6 s to respond. Trials were separated by a jittered inter-trial interval (2–3 s).

**Figure 1.**
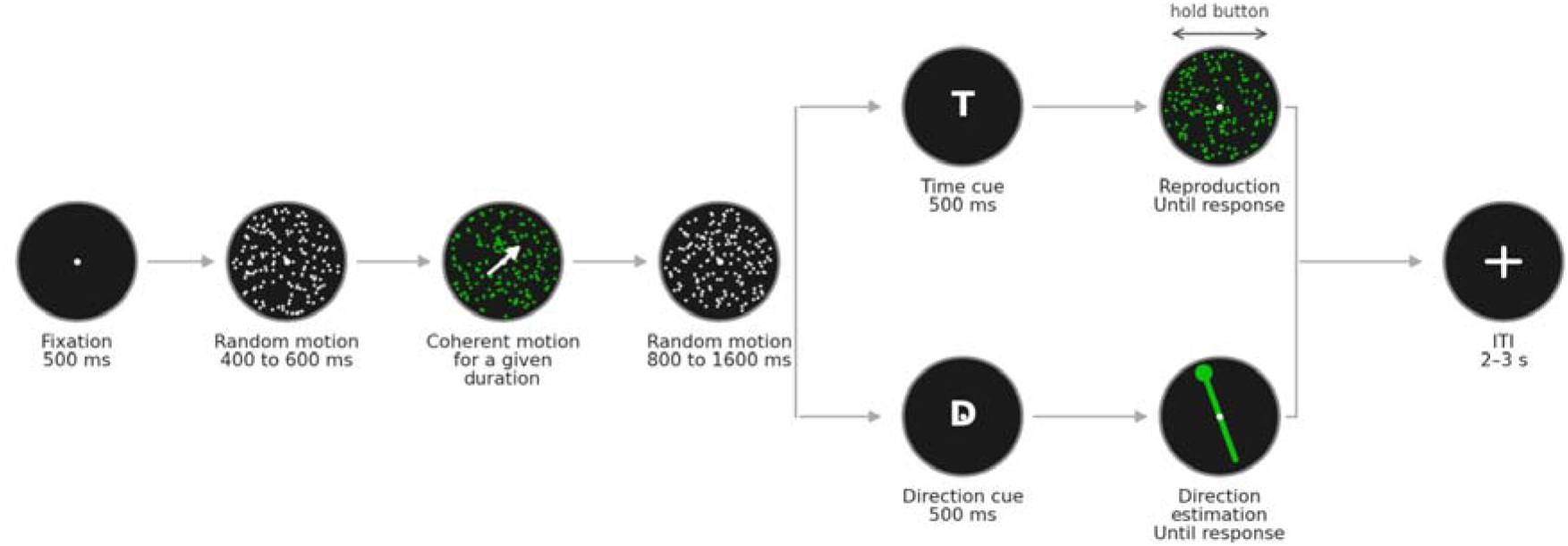
Illustration of a single trial. Each trial began with a fixation cross (0.5 s), followed by a random-motion mask (0.4–0.6 s). A coherent motion stimulus then appeared for its target duration (0.8–1.8 s), followed by a second random-motion mask (0.8–1.6 s). A retro-cue ("T" or "D") appeared for 0.5 s, indicating which feature to reproduce. For the time task ("T"), a green RDK display appeared; participants pressed a button to initiate the motion, held it for the perceived duration, and released it. For the direction task ("D"), a display with an adjustable line appeared; participants held the button to rotate the line toward the perceived motion direction and released it when satisfied. After their response, a 500-ms feedback display indicated accuracy. The next trial began after a jittered inter-trial interval (2–3 s).

For duration reproduction, participants pressed and held a button on an MRI-compatible response box; the duration of the button press constituted the reproduced interval. For direction reproduction, a response line appeared at a random offset from the target direction; while the button was held, the line rotated at 60°/s toward the target, and participants released the button when satisfied with the alignment. This differed from prior direction/orientation studies using mouse movements for fine orientation tuning because of movement constraints during MRI scanning. Additionally, we use button presses for both tasks to minimize differential motor effects on brain activity.

The durations of the two random-motion masks varied randomly from trial to trial, so the interval from trial onset to the retro-cue was not fixed; it averaged approximately 3 s (*M* = 3.02, *SD* = 0.42). All timing used in the models was taken from the frame-accurate presentation log (see Section 2.7.1).

Trial-to-trial transitions defined four conditions: TT (task repeat, time), DT (task switch to time), DD (task repeat, direction), and TD (task switch to direction), each occurring with approximately equal probability. The experiment comprised two runs, each containing 6 blocks of 40 trials (240 trials per run; 480 formal trials total).

### 2.4 Behavioral Analysis

The first trial of each block was excluded from all serial dependence analyses. Trials were further excluded if the participant made no response, if reproduction error fell more than 3 *SD* from that participant’s mean (duration) or exceeded 45° (direction), or if response latency fell more than 3 *SD* from that participant’s mean. Together, these criteria removed 3.2% of formal trials, most often for an outlying response latency. The same trial list was used for the imaging analyses, so excluded trials are absent from the event files and were not modeled in the GLM.

For duration, reproduction error was regressed on the previous duration (demeaned), separately for task-repeat (TT) and task-switch (DT) trials; the regression slope quantifies serial dependence strength. For direction, reproduction error was modeled using a Derivative of von Mises (DoVM) kernel (Kondo et al., 2022; Little and Clifford, 2025; Sadil et al., 2024), separately for DD and TD conditions:

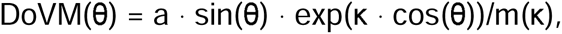

where θ is the angular difference between the previous and current direction, wrapped to ±180°, κ is the concentration parameter, and m(κ) is the peak of the bracketed kernel. Dividing by *m*(κ) sets the amplitude (*a*) equal to the peak of the fitted curve, so *a* is read directly in degrees of bias. A positive *a* indicates attraction toward the previous trial’s direction, and a negative repulsion from it.

Because curve height depends on *a* and κ jointly, the two trade off, and per-participant estimates of κ are poorly constrained when the underlying bias is small. We therefore estimated κ once from all direction trials pooled across participants, fitting separate amplitudes for DD and TD with a single shared κ so that their opposite-signed effects could not cancel (κ = 1.14, peak at 49°), and then held κ fixed while fitting each participant’s amplitude. Amplitudes were tested against zero with two-sided one-sample *t*-tests and, following Fischer and Whitney (2014), with a permutation test that shuffled the previous-trial direction within participant and condition across 5,000 iterations. Two robustness checks are reported: sensitivity to the fixed κ, and a fit-free measure defined as the mean signed error on trials where the previous direction led the current one minus trials where it lagged, computed within 90°.

### 2.5 Image Acquisition

MRI data were acquired on a 3T Siemens MAGNETOM Prisma scanner with a 32-channel head coil. Functional images used a T2*-weighted EPI sequence with multiband acceleration: TR = 1000 ms; TE = 30 ms; flip angle = 45°; FOV = 210 × 210 mm²; matrix = 70 × 70; voxel size = 3 × 3 × 3 mm³; 48 slices (no gap); multiband factor = 4; partial Fourier = 7/8. Each functional run comprised 2251 volumes (∼37.5 min). High-resolution T1-weighted structural images were acquired using an MPRAGE sequence (TR = 2500 ms; TE = 2.22 ms; voxel size = 0.8 × 0.8 × 0.8 mm³).

### 2.6 Image Preprocessing

Preprocessing was performed using *fMRIPrep* 25.1.2 (Esteban et al., 2020). Structural images were skull-stripped, tissue-segmented, and normalized to MNI152NLin2009cAsym space. Functional images underwent motion correction, slice-timing correction, co-registration, and resampling to 3-mm isotropic MNI space, matching the acquired resolution. BOLD time series were converted to percent signal change (PSC) units at this stage.

### 2.7 Statistical Analysis

#### 2.7.1 First-level general linear model

First-level analyses were performed using Nilearn 0.13. BOLD data were smoothed with a 6-mm FWHM kernel. Model fitting applied an additional constant rescaling, so unstandardized parameter estimates are reported in arbitrary units. A GLM was fitted to each participant’s concatenated two-run data with eight task regressors and six motion nuisance regressors. Event onsets were taken from the frame-accurate presentation log: retro-cue-locked regressors at each trial’s logged retro-cue onset, and the encoding boxcar spanning each trial’s onset-to-cue window. Because that window covaries with the stimulus duration (*r* = .68), the boxcar itself carries duration information, so the current-duration parametric modulator estimates duration-dependent variance over and above the boxcar (Section 3.2).

##### Stimulus-locked regressors

Two regressors captured stimulus-related activity: (1) a stimulus-onset boxcar spanning each trial’s full presentation window, from trial onset to retro-cue onset; (2) a current-duration parametric modulator (stick function at stimulus onset, modulated by Duration - 1.3 s), indexing regions whose onset-locked response scales with the duration of the coherent-motion stimulus.

##### Retro-*Cue-locked regressors* (onset = each trial’s logged retro-cue onset)

Four condition regressors modeled retro-cue-related activity as 0.5-s boxcar functions for TT, DT, DD, and TD transitions. Two parametric modulators captured serial dependence: a prior-duration modulator (previous trial’s duration, demeaned) applied to all time-cued trials (TT + DT), and a prior direction-difference modulator (angular difference wrapped to ±90°) applied to all direction-cued trials (DD + TD). Parametric modulators were pooled across transition types to maximize statistical power. A supplementary model splitting the prior-duration modulator into separate task-repeat and task-switch regressors is reported in the Supplementary Materials. The first trial of each block has no preceding trial, so it contributes to neither the condition regressors nor the modulators, while still being modeled by the stimulus-locked regressors above. Nilearn’s first-level GLM does not apply serial orthogonalization; all regressors were entered as separate columns, ensuring that each parameter estimate reflected only its unique contribution. All regressors were convolved with the canonical (Glover) HRF plus temporal derivative. The derivative terms were included to absorb small departures from the assumed response shape, but all contrasts weight the canonical column alone and place zero weight on the derivative; reported estimates are therefore canonical amplitudes. High-pass filtering (1/128 Hz) removed slow drifts, and temporal autocorrelation was modeled with an AR(1) process. Nuisance regressors comprised six rigid-body motion parameters. Full contrast definitions are provided in Supplementary Table S2, and the model specification itself is available in the public repository (see Data Availability).

#### 2.7.2 Control models for motor carryover

The primary model places 0.5-s boxcars at the retro-cue and contains no response regressors, so press-related activity can contribute to the transition coefficients. Conditions within each contrast share the same current task, but switch costs shift response onset. Because the two tasks differ substantially in hold duration, both the current and preceding responses are candidate sources. We therefore refitted the first-level model in two further variants, each adding regressors to the specification above and changing nothing else.

##### Response model

Two variable-duration boxcars, one per task domain, spanning each button press from key press to release. Responses without valid recorded press and release times were not included. These regressors model the variance associated with current and preceding press events. Because some response events were unavailable and the response and retro-cue regressors remain temporally correlated, the model cannot guarantee complete separation and is treated as a sensitivity analysis rather than a replacement for the primary model.

##### Previous-hold model

For each current-task domain, a retro-cue-locked stick regressor was parametrically modulated by the preceding trial’s hold duration. Previous trials were defined within block, and available hold values were mean-centred across trials of that current-task domain within each run. The model estimated the transition contrasts and prior-stimulus-duration coefficient conditional on this linear previous-hold covariate.

To verify that the added covariate captured motor-related variance, we conducted an exploratory positive-control ROI analysis of its canonical-HRF coefficient for current time-cued trials. Mean coefficients were extracted from fixed 8-mm spheres in left precentral gyrus (MNI -38, -22, 56; center labeled Precentral Gyrus by the Harvard–Oxford cortical atlas), SMA/pre-SMA (0, 10, 50), and right putamen (24, 4, 4). The latter two were the a priori timing ROIs defined below. Coefficients were tested against zero with two-sided one-sample *t*-tests across participants, with Benjamini– Hochberg FDR correction across the three diagnostic ROIs. These tests assess sensitivity of the covariate to motor-related variance; stability of the DT > TT contrast was assessed separately from its corrected extent and whole-brain map similarity across model variants.

#### 2.7.3 Group-level analysis

Whole-brain and region-of-interest analyses answer different questions, and we report both for that reason. The whole-brain permutation test searches without a spatial prior and controls the family-wise error rate across the brain, so it can find effects we did not anticipate, at a cost in sensitivity. The a priori ROIs test a small number of pre-specified regions with correspondingly more power, giving a confirmatory test that does not depend on a cluster surviving whole-brain correction. Regions defined from the present data serve neither purpose and are flagged as such wherever they appear.

Group-level inference used FSL *randomize* (Winkler et al., 2014) with threshold-free cluster enhancement (TFCE; Smith and Nichols, 2009). For each contrast, the 27 individual effect-size maps were concatenated and tested against zero in a one-sample design using 5000 sign-flip permutations, restricted to the intersection of the participants’ brain masks. Positive and negative directions were tested separately and both were reported, giving two-tailed inference at *p*_FWE_ < .05.

#### 2.7.4 Region-of-interest analysis

Eleven a priori ROIs were defined as 8-mm radius spheres centered on coordinates from the neuroimaging timing and task-switching literatures (Brass and von Cramon, 2002; Derrfuss et al., 2005; Wiener et al., 2010). Seven *timing-related* ROIs operationalized the predictions set out in the Introduction. SMA/pre-SMA (0, 10, 50), right putamen (24, 4, 4) targeted the cortical and striatal nodes of the STC circuit; right superior temporal gyrus (60, -20, 6) and right intraparietal sulcus (32, -52, 48) targeted the wider timing network identified by meta-analysis (Wiener et al., 2010); and bilateral MT/V5 (±45, -70, 0) and V1 (0, -84, 4) targeted the visual areas that represent the stimulus, the candidate substrate for direction carryover and the sites where the temporal-trace account predicts no prior-duration modulation. No sphere was placed on the thalamic node of the circuit: the timing-relevant thalamic territory has no consensus coordinate, and an 8-mm sphere at an atlas landmark would average across functionally distinct nuclei, so the thalamus was assessed by the whole-brain analysis, which retains full coverage of the structure. Four *executive-control* ROIs targeted regions implicated in task-switching (Brass and von Cramon, 2002): left dorsolateral prefrontal cortex (-42, 30, 26), dorsal anterior cingulate cortex (0, 24, 36), pre-SMA (0, 10, 50), and left inferior frontal junction (-44, 4, 30). Sphere centers are atlas landmarks (Harvard–Oxford, Juelich), except the left inferior frontal junction, which is a published coordinate (Brass and von Cramon, 2002). The medial premotor sphere (0, 10, 50) serves both families and is corrected separately for each. For each ROI, mean parameter estimates were extracted and tested with one-sample *t*-tests (FDR-corrected, *q* < .05, separately within each ROI family). To test whether task-switching effects differed by domain, we compared each participant’s Direction TD > DD and Time DT > TT estimates with two-sided paired *t*-tests in the four executive-control ROIs; Benjamini–Hochberg FDR correction was applied across these four comparisons, and paired effect sizes are reported as Cohen’s *d_z_*. Brain–behavior correlations between ROI parameter estimates and behavioral serial dependence measures were computed across all ROIs within each analysis family.

## 3. Results

### 3.1 Behavioral Serial Dependence

Duration reproduction exhibited robust attractive serial dependence: current reproductions were systematically biased toward the previous trial’s duration, as indicated by the regression analysis (Figure 2A). Serial dependence was approximately four times stronger on task-repeat (TT: mean slope = 0.13, SD = 0.07) than task-switch (DT: mean slope = 0.03, SD = 0.06) trials, *F*(1, 26) = 22.49, *p* < .001, η²_p_ = .46, *d* = 0.91. This pattern replicates Cheng et al. (2024b) and confirms that temporal serial dependence is amplified when the same task is repeated across consecutive trials.

**Figure 2.**
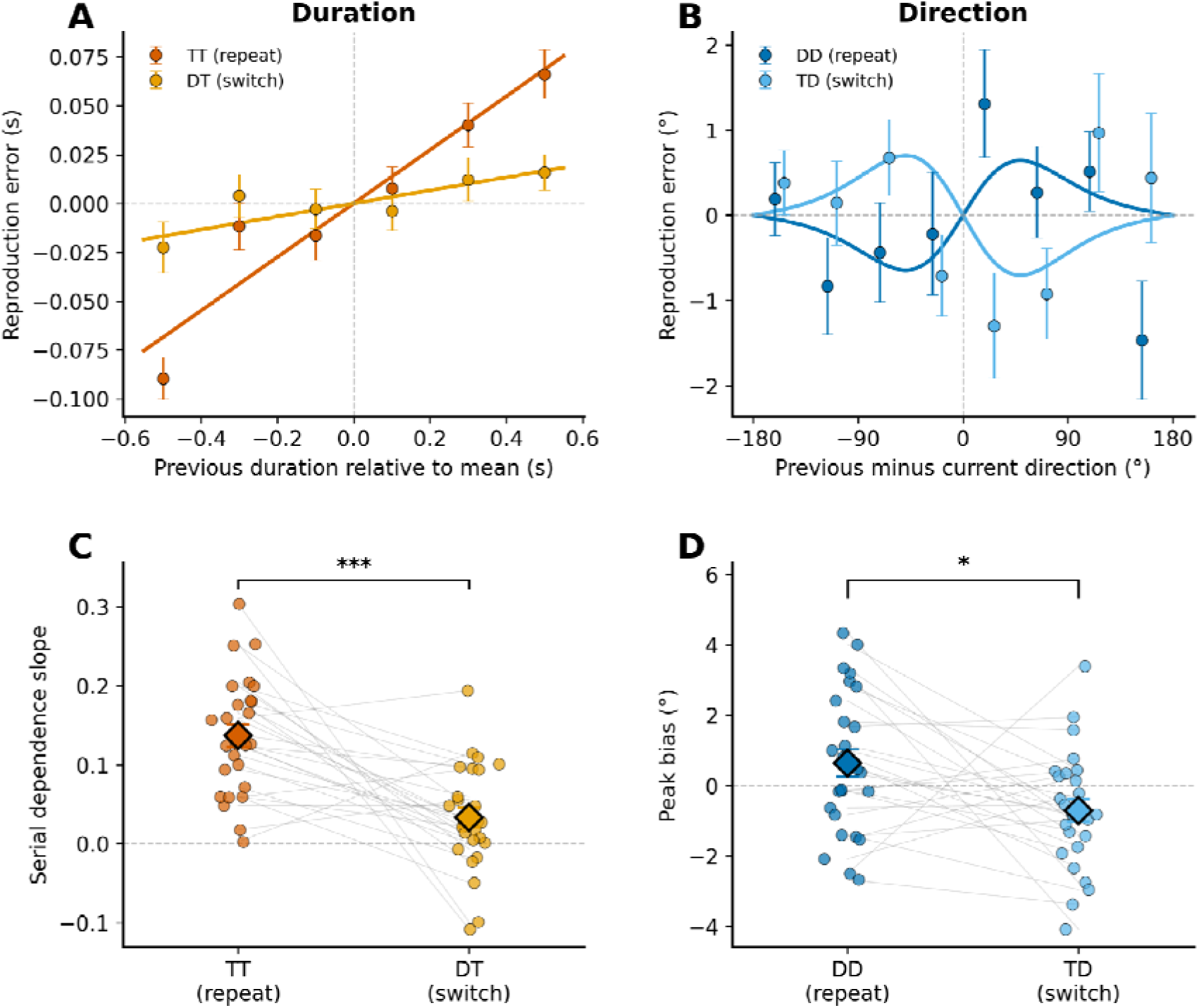
Behavioral serial dependence (*N* = 27). (**A**) Duration reproduction error against the previous trial’s duration, expressed relative to the 1.3-s grand mean, for task-repeat (TT) and task-switch (DT) trials. (**B**) Direction reproduction error against the angular difference between the previous and current direction, wrapped to ±180° and binned at 45°, for task-repeat (DD) and task-switch (TD) trials; the two conditions are offset horizontally by 4° for legibility. In (**A**) and (**B**), points are group means with SEM, obtained by averaging within each participant before averaging across them, and error was demeaned within participant and condition. The overlaid line and curve are the group fits reported in the text, not refits of the binned means, so the points give an independent view of how well each function describes the data. (**C**) Duration serial dependence slopes and (**D**) direction serial dependence as the peak of the fitted DoVM curve in degrees, where positive values denote attraction toward the previous direction and negative values repulsion. Dots = individual participants; diamonds = group mean ± SEM. *** *p* < .001, * *p* < .05.

Direction reproduction carried a much smaller bias that changed sign with the preceding task (Figure 2B). Reproductions were drawn toward the previous direction when the direction task repeated (DD: peak bias = +0.64°, 95% CI [-0.14, 1.43], *d_z_* = 0.32, *p* = .105, permutation *p* = .059) and pushed away from it when the task switched (TD: peak bias = -0.70°, 95% CI [-1.36, -0.05], *d_z_* = -0.42, *p* = .036, permutation *p* = .034). Neither condition alone is decisive, but they differ reliably, DD - TD = 1.35°, 95% CI [0.25, 2.44], *F*(1, 26) = 6.39, *p* = .018, η²_p_ = .20, permutation *p* = .003. The fit-free measure reproduces this pattern (DD = +0.96°, *p* = .176; TD = -1.22°, *p* = .032; difference = 2.18°, *p* = .022), and the difference holds for any fixed κ between 0.5 and 3.

It should be noted that the direction biases are of the order of one degree, about 6% of the trial-to-trial spread in direction reproduction error, and the button-press rotation method is coarser than the continuous adjustment used elsewhere (Cheng et al.,2024b), so these estimates are conservative. The duration effect is roughly four times larger, reaching about 24% of the spread in duration reproduction error. The two features of the same stimulus therefore carry histories that differ in size and in sign, which is what a domain-specific account predicts.

Response onset latency revealed a significant switch cost in the time domain: DT trials were 39 ms slower than TT trials, *t*(26) = 3.72, *p* = .001, *d* = 0.72. Direction showed a smaller, non-significant switch cost (8 ms, *t*(26) = 1.87, *p* = .073, *d* = 0.36). The Domain × Switch interaction confirmed this asymmetry, *t*(26) = 2.84, *p* = .009, *d* = 0.55. Reproduction accuracy did not differ between switch and repeat trials in either domain (both *p*s > .28), indicating that the switch cost was expressed in response initiation speed, not reproduction precision.

### 3.2 Current Duration Encoding

To validate the paradigm’s sensitivity to temporal processing, we examined the current-duration parametric modulator. Whole-brain TFCE analysis revealed widespread positive modulation (*p*_FWE_ < .05) spanning 5.7% of the analyzed volume. The largest cluster (62,127 mm³; peak *t* = 8.28 at MNI -24, -42, -9) is a single bilateral posterior swath: its peak lies in left occipitotemporal cortex, but it extends across occipital, temporal, and inferior parietal cortex in both hemispheres. Two further clusters centered on right insula/putamen (11,151 mm³; MNI: 33, 6, 12) and left insula (7,236 mm³; MNI: -42, -9, 0), regions repeatedly implicated in duration estimation (Coull et al., 2011; Wittmann et al., 2010). Smaller negative clusters were confined to left frontal operculum (4,212 mm³) and medial premotor cortex (864 mm³). ROI analysis showed duration-dependent modulation in SMA/pre-SMA, *t*(26) = -3.90, *d* = -0.74, *p* = .0069, regions involved in timing (Baykan et al., 2026 ; Wiener et al., 2010).

It should be noted that the encoding boxcar spans each trial’s presentation window, and that window covaries with the stimulus duration (*r* = .68), so the boxcar itself carries duration information. The parametric modulator, therefore, indexes duration-dependent residual scaling over and above the boxcar, and its sign should be read accordingly. What the analysis establishes for present purposes is that the paradigm engages duration-sensitive responses across timing-related cortex and subcortex, providing the baseline against which serial dependence effects can be evaluated (see Supplementary Figure S1).

### 3.3 Neural Correlates of Serial Dependence

#### 3.3.1 Duration: a prior-duration signal in the putamen

The duration of the prior trial modulated the current retro-cue-locked neural activity in the bilateral putamen, the region to survive whole-brain correction (Figure 3).

**Figure 3.**
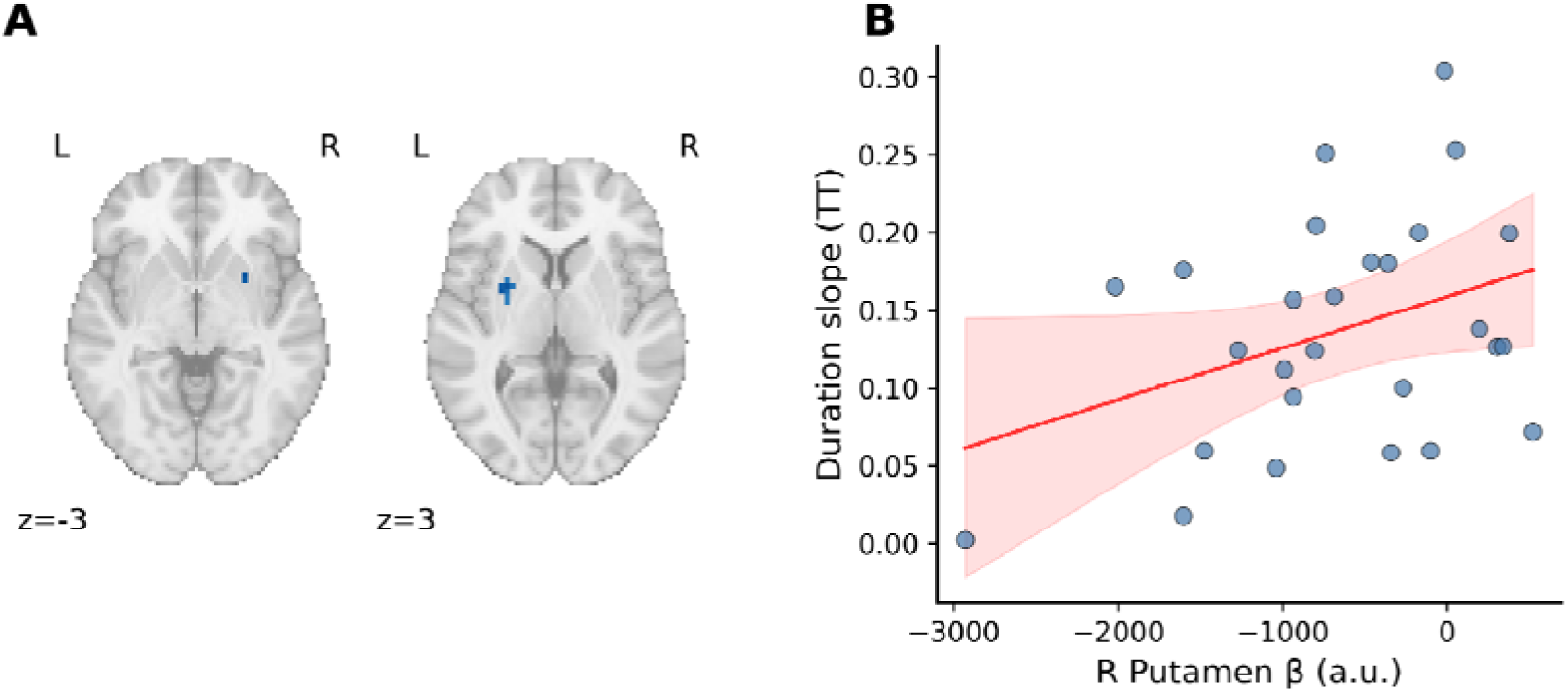
Prior-duration modulation. (**A**) Negative prior-duration modulation, thresholded using voxelwise TFCE-FWE correction at p < .05 and shown on axial sections at z = −3 and 3. Blue denotes more negative signed t values. Significant components were observed in the left putamen (540 mm³; peak MNI: −30, 0, 0; peak t = −6.32, p_FWE_ = .021) and right putamen (81 mm³; peak MNI: 27, 9, −3; peak *t* = −6.01,p_FWE_= .041). (**B**) Right putamen parameter estimate against behavioral duration serial dependence on task-repeat trials, one point per participant, with least-squares fit and 95% confidence band (*r* = .37, 95% CI [-0.02, 0.66], *p* = .060). Parameter estimates are extracted from an anatomical ROI independent of the whole-brain components in panel A.

Both surviving clusters of TFCE-FWE-significant voxels localized to the putamen, and both were negative, longer preceding durations producing weaker responses: left (20 3-mm isotropic voxels, 540 mm³; MNI: -30, 0, 0; signed peak *t* = -6.32, *p*_FWE_ = .021) and right (3 voxels, 81 mm³; MNI: 27, 9, -3; signed peak *t* = -6.01, *p*_FWE_ = .041). The right putamen ROI, specified a priori from the timing literature (Section 2.7.4), confirmed the effect, *t*(26) = -4.20, *d* = -0.81, *p*_FDR_ = .005.

No cluster reached significance in the positive direction. Because the strongest positive voxels lay in the posterior medial cortex, regions our earlier analysis had highlighted, we tested those regions directly under two definitions. In 8-mm spheres centered on the positive peaks, the effects looked substantial: posterior cingulate cortex (PCC, *d* = +0.57, *p* = .006), precuneus (*d* = +0.49, *p* = .017), and occipital pole (*d* = +0.51, *p* = .014). Those centers derive from the data themselves. Repeating the test in anatomically defined Harvard–Oxford parcels, chosen without reference to any statistical map, left none of them reliable: PCC *d* = +0.12 (*p* = .54), precuneus *d* = +0.38 (*p* = .06), occipital pole *d* = +0.11 (*p* = .56). The gap between the two tests measures what selection at the peak buys. Prior-duration modulation that survives both whole-brain correction and independent regional definition is confined to the putamen.

None of the remaining a priori ROIs showed modulation: SMA/pre-SMA, right STG, bilateral MT/V5, right IPS, and V1 all gave |*d*| < 0.07 (all *p*_FDR_ = .99). The nulls in V1 and MT/V5 locate this result within the wider serial dependence literature.

For orientation and motion, the carryover signal has a demonstrated locus in early visual cortex, where repulsive population shifts accompany an attractive behavioral bias (John-Saaltink et al., 2016; Sheehan and Serences, 2022). A prior-duration signal appears instead in the dorsal striatum, the structure the timing literature identifies as encoding elapsed intervals (Coull et al., 2011; Merchant et al., 2013), and in no sensory region we tested. The brain–behavior correlation between right putamen modulation and behavioral serial dependence was only marginally significant (*r* = .37, 95% CI [-0.02, 0.66], *p* = .060, uncorrected; Figure 3B). A supplementary condition-separated analysis showed numerically stronger putamen suppression on task-repeat than task-switch trials, though the direct comparison did not reach significance (Supplementary Table S1).

Because the primary logged-timing GLM did not include response regressors, we tested robustness to the 423-ms longer preceding hold before DT than TT. Adding variable-duration response boxcars left the putamen estimate essentially unchanged (*d* = -0.85; per-subject *r* = .99 with the primary model), as did entering preceding hold duration as a competing retro-cue-locked covariate (*d* = -0.79; *r* = .99). In the positive-control analysis, that covariate showed negative effects in left precentral gyrus, *t*(26) = - 2.44, *d* = -0.47, *p* = .022, *q*_FDR_ = .033, and SMA/pre-SMA, *t*(26) = -2.98, *d* = -0.57, *p* = .006, *q*_FDR_ = .018, but not right putamen, *t*(26) = -0.46, *d* = -0.09, *p* = .65, *q*_FDR_ = .65 (Supplementary Table S4). Thus, the covariate captured variance in motor regions while the putamen prior-duration estimate remained stable. These sensitivities argue against measured hold-duration carryover as the sole explanation but do not establish that the putamen effect is motor-independent.

#### 3.3.2 Direction: Null neural serial dependence

The prior direction-difference modulator produced no significant clusters at the whole-brain level. All timing-related ROIs showed negligible modulation (|*d*| < 0.21, all *p*_FDR_ > .60), including MT/V5, IPS, and V1. Thus, the detected prior-duration effect had no counterpart in the direction domain in this paradigm. Because the modulators differ and the behavioral direction effect was weak, this null does not establish a domain-specific neural architecture.

### 3.4 Neural Correlates of Task-Switching

#### 3.4.1 Time domain: Asymmetric switch cost

Switching from direction to time (DT > TT) activated 553,743 mm³, 39.1% of the analyzed volume in the primary logged-timing model (Figure 4A). ROI analysis showed left IFJ as the strongest executive-control hub, *t*(26) = 5.35, *d* = 1.03, *p*_FDR_ < .001, followed by left DLPFC, *d* = 0.71, *p*_FDR_ = .003. Pre-SMA (*d* = 0.38) and dACC (*d* = 0.03) did not reach significance. The reverse contrast (TT > DT) produced no significant voxels.

**Figure 4.**
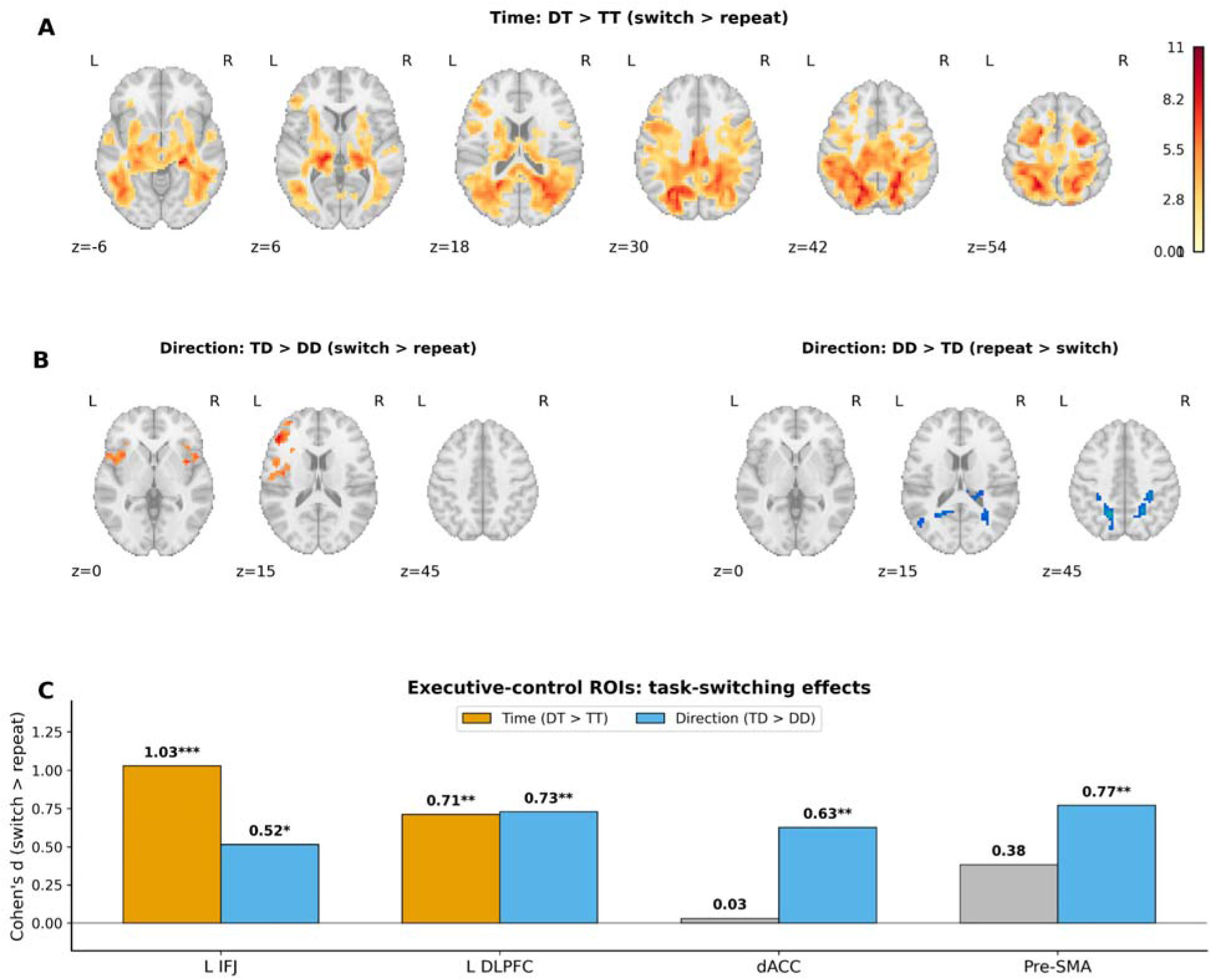
Task-switching contrasts from the primary logged-timing model (*p*_FWE_ < .05). (**A**) DT > TT, 553,743 mm³. (**B**) TD > DD (left) and DD > TD (right). (**C**) Executive-control ROI effect sizes for both domains. * *p*_FDR_ < .05; ** *p*_FDR_ < .01; *** *p*_FDR_ < .001.

DT responses were 39 ms slower than TT responses, but current hold duration and the presentation window were balanced. The widespread DT > TT pattern remained when each press-to-release response was modeled explicitly (613,035 mm³, 43.3% of the analyzed volume; spatial correlation with the primary map, *r* = .966) and when preceding hold duration was entered as a competing retro-cue-locked modulator (444,042 mm³, 31.3%; *r* = .969). The previous-hold covariate itself showed FDR-corrected effects in left precentral and SMA/pre-SMA (Section 3.3.1; Supplementary Table S4), confirming sensitivity to motor-related variance even though the DT > TT pattern remained spatially stable. These analyses support stability to the modeled response-timing variables. They do not isolate a purely attentional or reconfiguration process because DT > TT necessarily differs in the preceding task domain, and the explicit response model has elevated collinearity (median Time-DT VIF = 10.48; maximum = 20.85; Supplementary Figure S2 and Tables S3-S4).

#### 3.4.2 Direction-domain transition effects

Switching from time to direction (TD > DD) activated 18,576 mm³, with peaks in left (MNI: -33, 21, 9) and right (36, 27, 9) frontal operculum and in SMA/pre-SMA (6, 3, 66), and engaged all four executive-control ROIs (all *p*_FDR_ < .05; Figure 4B). The dACC showed a reliable direction-switching effect (*d* = 0.63), whereas its time-switching estimate was near zero (*d* = 0.03). A direct paired comparison suggested a larger dACC effect for direction than time switching, *t*(26) = 2.37, *d_z_*= 0.46, *p* = .026, but the difference did not survive FDR correction across the four executive-control ROIs, *p*_FDR_ = .103. No other between-domain ROI comparison survived correction (all *p*_FDR_ ≥ .170). The dACC pattern is therefore suggestive rather than evidence of domain specificity. Repeating direction (DD > TD) activated a complementary sensorimotor network of 47,169 mm³ spanning superior parietal cortex (44,820 mm³; peak -21, -60, 57) and bilateral precentral gyrus (Figure 4B).

#### 3.4.3 Separability of the two signals

The thresholded prior-duration and task-transition maps occupy different territory (Figure 5). Reliable prior-duration modulation was detected in the putamen, where no transition contrast was significant; the transition contrasts recruited frontoparietal cortex (IFJ, DLPFC, dACC, pre-SMA), where no prior-duration effect was significant. The retro-cue design allows previous-duration modulation and retro-cue-locked transition effects to be estimated as distinct regressors within the same participants rather than inferred across studies. Non-overlap among thresholded effects does not, however, constitute a formal dissociation or identify the cognitive process responsible for the transition contrast.

**Figure 5.**
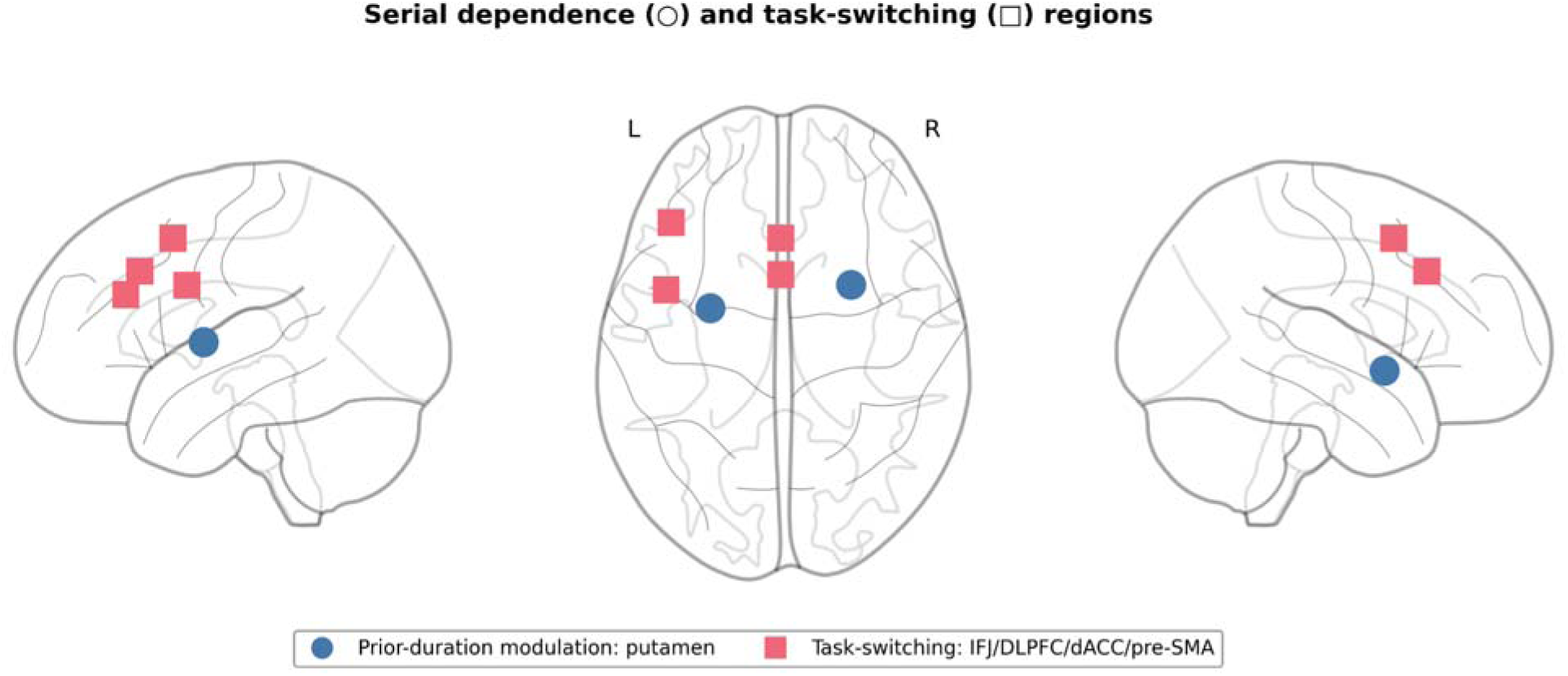
Anatomical summary. Circles mark the peaks of the two prior-duration clusters (bilateral putamen); squares mark the a priori executive-control ROIs engaged by task switching (IFJ, DLPFC, dACC, pre-SMA). The two sets do not overlap.

Supplementary analyses (condition-separated parametric modulators and additional whole-brain maps) are reported in the Supplementary Materials.

## 4. Discussion

This study identified a retro-cue-locked neural correlate of previous duration in the bilateral putamen. Longer preceding durations predicted weaker putamen activity, and this negative modulation was the only prior-stimulus effect to survive whole-brain correction. Analysis of the a priori putamen region of interest, defined from the timing literature, confirmed the result. The finding narrows our original prediction that duration serial dependence would engage the wider striato-thalamo-cortical circuit: the circuit’s cortical node, SMA, showed no reliable prior-duration modulation, and the thalamus showed no effect at the whole-brain level. The wider timing network showed no better, with STG and IPS both null. Temporal carryover, therefore, appeared most clearly in the striatal component of the timing network, although the regional pattern does not identify a particular processing stage within the basal ganglia loop (Meck et al., 2008; Yin et al., 2022).

Locating this signal in the putamen extends the earlier report of prior-duration information in the caudate (Cheng et al., 2024a). The two structures occupy different positions within the dorsal striatum: the caudate receives predominantly associative and prefrontal input, whereas the putamen is more closely linked to sensorimotor circuits (Grahn et al., 2008). Yet the putamen’s function is not confined to movement. Basal ganglia disorders can affect memory and decisions about time, even when the task requires no timed movement, placing striatal function within temporal cognition as well as motor control (Allman and Meck, 2012; Wearden et al., 2008). That clinical evidence does not isolate the putamen, since these conditions affect the striatum broadly, but our response-control analyses point in the same direction. The prior-duration estimate remained stable after modeling current responses and preceding hold duration, while the hold-duration covariate captured variance in left precentral and medial premotor cortex. This pattern argues against measured hold duration as the sole source of the putamen effect, without excluding contributions from other motor or sensory history. Because the present and earlier studies also differ in task, scanner, sample, and analysis pipeline, the caudate-putamen contrast is best treated as a hypothesis that task demands influence where prior-duration signals emerge within dorsal striatum.

The negative sign of the modulation constrains, but does not determine, its mechanism. Reduced retro-cue-locked BOLD after a longer preceding duration could reflect active attenuation of retained information, altered dopaminergic state after timing a longer interval, or adaptation within the machinery that accumulates temporal evidence. These accounts all predict a relationship with the previous duration, but they differ in what that relationship represents. In particular, a negative coefficient does not by itself demonstrate active suppression of a stored trace. The inconclusive brain-behavior association likewise leaves open whether the putamen modulation contributes directly to the attractive behavioral bias or reflects another consequence of the preceding timing episode.

No reliable positive prior-duration modulation remained outside the striatum once posterior medial regions were defined independently of the statistical map. Peak-centered spheres suggested sizeable PCC and precuneus effects, but anatomically defined parcels did not reproduce them. Posterior medial cortex has nevertheless been implicated in integrating temporal context across distributional learning and passive encoding (Baykan et al., 2026 ; Cheng et al., 2024a). Whether it carries trial-to-trial duration information remains open and requires independently specified parcels, a functional localizer, or preregistered coordinates.

Motion direction behaved differently from duration in both measures. Behaviorally, it produced a bias of about one degree that reversed sign with the preceding task, attractive when the direction task repeated and repulsive when it switched, against a duration effect roughly four times larger in the same participants once each is scaled to its own error spread. The repulsion on switch trials is consistent with the previous finding (Bae and Luck, 2020). Neurally, there was no counterpart to this small effect. The neural null cannot distinguish absent directional carryover from limited sensitivity to its retro-cue-locked expression. Work on orientation shows that attractive behavior can coexist with repulsive adaptation in the early visual cortex (John-Saaltink et al., 2016; Sheehan and Serences, 2022). Directional carryover may follow a similar sensory-adaptation-plus-readout architecture whose retro-cue-phase signature escaped detection here. The present within-participant comparison therefore identifies no direction-domain counterpart to the putamen signal, but it does not establish a domain-specific neural architecture.

Task transitions recruited a second, broader set of regions. Switching from direction to time produced widespread activation, strongest among the executive-control ROIs in left IFJ and DLPFC, consistent with the role of IFJ in updating task representations (Derrfuss et al., 2005). The spatial pattern held when observed responses and preceding hold duration each had their own regressors, making unequal response timing an unlikely sole explanation. The contrast nonetheless remains tied to the preceding task domain, and can combine executive control, attention, and pre-response preparation. Direction switching produced a more focal pattern that included dACC. A direct comparison suggested a stronger dACC response for direction than time switching, but the difference did not survive correction across the executive-control ROI family.

The thresholded prior-duration and transition maps implicated different regions: the putamen carried the prior-duration signal, whereas IFJ and DLPFC dominated the time-transition effect. This descriptive separation raises the possibility that a retained duration-related state and executive transition processes make distinct contributions to behavior, but it does not demonstrate an anatomical dissociation or a gating mechanism. Behavioral evidence provides a more specific rationale for testing gating. Temporal serial dependence weakens when the task changes, and Wu et al. (2026) found that it vanished when reproduction and bisection were interleaved under a post-cue design that held duration encoding constant while varying the motor response. A bound-event-file account could explain this dependence if duration and response mode are encoded together (Frings et al., 2020). The present condition-separated analysis did not detect a reliable difference in putamen modulation between repeat and switch trials. However, the design changed reported features, response mode, preceding task, and motor demands together. Response-mode gating, therefore, remains a plausible hypothesis rather than a result of this study.

Testing that hypothesis requires a design that crosses task domain and response mode while providing enough trials to estimate their interaction. Multivariate analyses could then ask whether retro-cue-phase putamen patterns retain information about the previous duration. Successful decoding would establish duration-related information in the pattern, but would not alone distinguish a stored representation from adaptation or a changed network state. That distinction requires model-specific predictions about pattern geometry, temporal evolution, or sensitivity to orthogonal manipulation. Patient, pharmacological, or stimulation studies could further test whether changing putamen function alters temporal serial dependence, provided that general timing and motor performance are measured separately.

Several limitations set the scope of these conclusions. The primary parametric modulator pooled task-repeat and task-switch trials, and the lower-powered condition-separated model could not determine whether putamen modulation differs between them. The DT > TT contrast is inseparable from the preceding task domain; response-control models address measured response timing, not every motor, sensory-history, or attentional consequence of that difference. Previous stimulus duration is correlated with the previous visual-exposure duration, so the prior-duration coefficient cannot distinguish a mnemonic signal from exposure-dependent adaptation. Fixed-radius ROIs do not accommodate individual anatomy, making null results sensitive to the choice of coordinates. Finally, the button-press direction method was tailored to MRI scanning and may have obscured subtle repulsive biases that continuous adjustment can reveal.

In conclusion, the present study establishes a focal subcortical correlate of duration serial dependence: previous duration modulated bilateral putamen activity, whereas no reliable counterpart emerged in the sensory regions or direction task tested here. This pattern differs from the early visual-cortical signatures reported for orientation and motion, motivating the hypothesis that temporal history is expressed through a different neural route. Task transitions engaged frontoparietal control regions, but the present data do not show that these regions regulate the putamen signal. The broader implication is that serial dependence should not be treated as a unitary neural operation inferred from similar behavioral biases across domains. A neural account must explain both what information from the recent past is retained and how current task demands alter its influence. The putamen finding anchors the first question for duration, while the unresolved relation between striatal carryover and task-transition activity defines the evidence needed to answer the second. Deciding whether switching weakens the trace itself or gates its readout requires showing that frontoparietal transition activity conditions the striatal signal: for example, trial-wise coupling between IFJ and putamen that strengthens on switch trials and predicts attenuation of the prior-duration slope. Such a test demands more trials per condition than the present design provides.

## CRediT Author Contribution Statement

**Si Cheng:** Conceptualization, Data curation, Formal Analysis, Investigation, Methodology, Writing, Review, and Editing. **Siyi Chen:** Investigation, Funding acquisition, Writing, Review, and Editing. **Jiao Wu**: Data Curation. **Chunyu Qu**: Formal Analysis, Investigation, Methodology. **Zhuanghua Shi**: Conceptualization, Funding acquisition, Project administration, Supervision, Writing, Visualization, Review, and Editing.

## Declaration of Competing Interests

The authors declare no competing interests.

## Funding

This work was supported by German Research Foundation DFG grants CH3093/1-1 and SH 166/10-1, awarded to Siyi Chen and Zhuanghua Shi, respectively, and was carried out using the NICUM Siemens Prisma scanner supported by DFG grant INST 86/1739-1 FUGG.

## Data Availability

The raw MRI and behavioral data, the frame-accurate per-trial timing recovered from the presentation logs, and the analysis code are deposited on G-Node Infrastructure at https://gin.g-node.org/msense/time-direction-fmri-dataset.

## Declaration of Generative AI Use

During the preparation of this work, the authors used Claude AI and Grammarly AI to polish the language and tidy and optimize Python analysis codes. After using these tools/services, the authors reviewed and edited the content as needed and take full responsibility for the published article.

## Notes

### Competing Interest Statement

The authors have declared no competing interest.

### Summary of Updates

During our revision, we found some deviation in our own stimulus presentation code. Each trial presented a random-motion mask, the coherent-motion stimulus, and a second random-motion mask before the retro-cue. The masks were designed to complement the stimulus so that the stimulus-to-cue interval remained fixed at 3 s, regardless of the duration being judged. The mask durations (before and after the stimulus) were generated correctly, but the trial list was shuffled column-wise, which broke their pairing with the corresponding durations. So we re-ran the first-level GLM with each trial's true cue onset, and applied the encoding boxcar to each trial's true stimulus-to-cue window, and the results and discussion were changed accordingly. Additionally, we open our MRI data at https://gin.g-node.org/msense/time-direction-fmri-dataset.

