## Supplementary for "Neural Substrates of Duration Serial Dependence: Putamen Suppression of the Prior Duration Trace"

### **1. Condition-Separated Prior-Duration Parametric Modulator**

The main GLM pooled the prior-duration parametric modulator across task-repeat (TT) and task-switch (DT) trials to maximize statistical power (~24 trials per run). To ask whether the putamen suppression was present on both trial types or was task-gated, a supplementary GLM split the prior-duration modulator into separate TT and DT regressors (~12 trials each per run). The model was otherwise identical to the main analysis, including the logged cue onsets and the presentation-duration encoding window.

Mean parameter estimates were extracted from four ROIs: one a priori, right putamen (24, 4, 4; 8-mm sphere), and three centered on whole-brain peaks — left putamen (-30, 0, 0), PCC (-3, -33, 27), and precuneus (6, -63, 48; all 8-mm spheres). The latter three are not independent of the data, and the two posterior medial coordinates come from a positive cluster that does not survive whole-brain correction in the present analysis; they are reported for completeness and not as independent tests.

**Table S1.** Condition-separated prior-duration modulator: ROI statistics.

| **ROI** | **TT mean (SEM)** | **TT  *d*** | **TT  *p*** | **DT mean (SEM)** | **DT  *d*** | **DT  *p*** | **TT vs DT *p*** |
| --- | --- | --- | --- | --- | --- | --- | --- |
| R Putamen (a priori) | -937 (249) | -0.73 | < .001 | -390 (245) | -0.31 | .123 | .154 |
| L Putamen† | -970 (257) | -0.73 | < .001 | -321 (247) | -0.25 | .206 | .134 |
| PCC† | -408 (325) | -0.24 | .219 | -461 (279) | -0.32 | .111 | .902 |
| Precuneus† | 619 (412) | 0.29 | .145 | -286 (396) | -0.14 | .476 | .178 |

*Note.* Mean values are unstandardized parameter estimates (arbitrary units). One-sample *t*-tests assessed whether estimates differed from zero; paired *t*-tests compared TT vs DT. *N* = 27. †Coordinate defined at a whole-brain peak and therefore not independent of the data.

Bilateral putamen suppression was reliable on task-repeat trials (right *d* = -0.73, left *d* = -0.73, both *p* < .001) and did not reach significance on task-switch trials (right *d* = -0.31, *p* = .123; left *d* = -0.25, *p* = .206). The direct TT versus DT comparisons were themselves not significant (right *p* = .154; left *p* = .134). With roughly twelve trials per condition per run this analysis is not powered to estimate the interaction, and we restrain our interpretation about whether the prior-duration trace differs between repeat and switch trials. Neither posterior medial ROI showed reliable modulation on either trial type.

### **2. First-Level Contrast Definitions**

**Table S2.** First-level contrasts in the primary general linear model.

| **Contrast** | **Interpretation** | **Nonzero weights on canonical-HRF columns** |
| --- | --- | --- |
| Current duration | Duration-dependent residual scaling at stimulus onset | +1 Current_duration |
| Time TT | Time-task repeat activity relative to the implicit baseline | +1 Time_TT |
| Time DT | Time-task switch activity relative to the implicit baseline | +1 Time_DT |
| Direction DD | Direction-task repeat activity relative to the implicit baseline | +1 Direction_DD |
| Direction TD | Direction-task switch activity relative to the implicit baseline | +1 Direction_TD |
| Time switch > repeat | Time-task switching effect | +1 Time_DT; -1 Time_TT |
| Time repeat > switch | Reverse Time-task switching effect | +1 Time_TT; -1 Time_DT |
| Direction switch > repeat | Direction-task switching effect | +1 Direction_TD;  -1 Direction_DD |
| Direction repeat > switch | Reverse Direction-task switching effect | +1 Direction_DD;  -1 Direction_TD |
| Prior duration | Prior-duration parametric modulation pooled across TT and DT trials | +1 nBack_duration |
| Prior direction difference | Prior direction-difference parametric modulation pooled across DD and TD trials | +1 nBack_direction_diff |

*Note.* The same contrast vector was applied to both runs. All unlisted task columns, six motion regressors, cosine drift terms, and the intercept received weight 0. Although each task regressor was modeled with a canonical Glover hemodynamic response function (HRF) and its temporal derivative, every contrast assigned weight 0 to the derivative column; reported estimates therefore reflect canonical-HRF amplitudes only. TT = Time-to-Time; DT = Direction-to-Time; DD = Direction-to-Direction; TD = Time-to-Direction.

### **3. Current-Duration Whole-Brain Map**

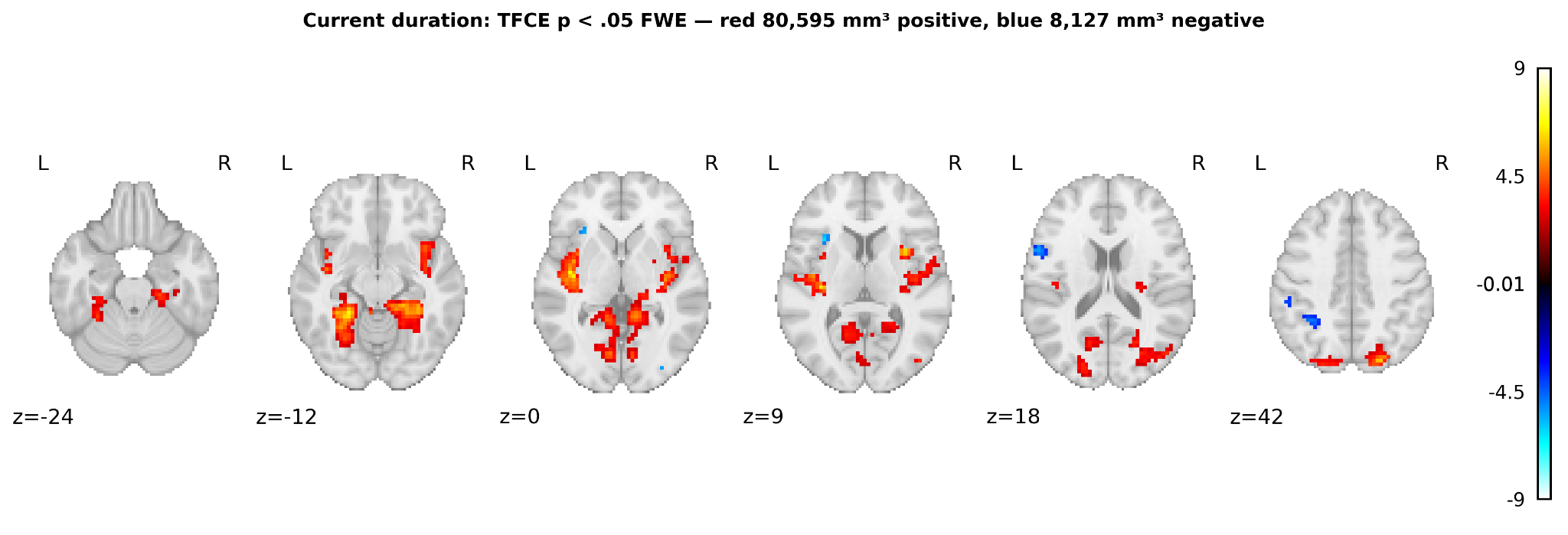

**Figure S1.** Current duration encoding: whole-brain TFCE maps (*p_FWE_* < .05), thresholded *t*-statistics on axial slices. Red, greater onset-locked BOLD for longer durations (80,595 mm³ across 5 clusters); blue, reduced BOLD for longer durations (8,127 mm³ across 9 clusters). The positive effect is dominated by a single bilateral posterior cluster (62,127 mm³; peak *t* = 8.28 at MNI -24, -42, -9) extending across occipital, temporal and inferior parietal cortex, with further clusters in right insula/putamen (11,151 mm³) and left insula (7,236 mm³). Negative clusters are confined to left frontal operculum (4,212 mm³; peak *t* = 9.10 at -57, 6, 24) and medial premotor cortex (864 mm³). As noted in Section 3.2, the encoding boxcar spans each trial's presentation window and therefore itself carries duration information, so this modulator indexes duration-dependent residual scaling rather than the total duration response.

### **4. Task-Switching Sensitivity Analyses**

The primary task-switching model uses 0.5-s cue-locked transition regressors and does not contain an explicit response regressor, so temporally correlated response activity can contribute to the transition coefficients. We therefore examined whether the widespread DT > TT activation was stable to differences in presentation timing, current response timing, or carryover from the preceding response.

#### 4.1 Event-timing diagnostics

DT responses began 39 ms later than TT responses, 95% CI [17, 61] ms, *t*(26) = 3.72, *p* = .001. This difference reflected response onset rather than press duration: current hold duration differed by less than 1 ms, 95% CI [−16, 14] ms, *p* = .916. The logged presentation window was also balanced, DT − TT = −6 ms, 95% CI [−26, 14] ms, *p* = .533. The preceding response was structurally different because DT follows a direction response, whereas TT follows a time response. On the trial preceding a DT trial, the button was held 423 ms longer before DT than on the trial preceding a TT trial, 95% CI [379, 467] ms, *p* < .001. This imbalance motivated the two response-control models below.

#### 4.2 Response-control models

Three alternative specifications were compared with the primary model. The nominal-timing model tested how much the result depends on frame-accurate event times: cue-locked regressors were placed at the nominal cue time (3 s after trial onset) and the encoding boxcar was held at a fixed 3-s duration, in place of the logged cue onsets and trial-specific presentation windows used throughout the main analyses. The response model added separate Time and Direction press-to-release boxcars for responses with valid recorded onsets and durations, modeling measured variance associated with current responses and retained preceding responses. It cannot guarantee complete separation because some preceding responses were unavailable and cue- and response-related regressors overlap. In the previous-hold model, each current-task domain received a cue-locked stick regressor modulated by the preceding trial's hold duration. Previous trials were identified within block, and available hold values were mean-centered across trials of that current-task domain within each run. The model, therefore, tested whether the transition pattern remained when this linear previous-hold covariate competed for variance. The response and previous-hold models otherwise retained the primary model's logged cue timing, trial-specific presentation boxcar, transition regressors, temporal derivatives, motion parameters, drift basis, and AR(1) correction.

**Table S3.** DT > TT sensitivity to timing and response-model specifications.

| **Model** | **Added control** | **Time-DT VIF, median (maximum)** | **Significant extent (mm³)** | **Analysis mask (%)** | **Map similarity, *r* / Dice** |
| --- | --- | --- | --- | --- | --- |
| Primary logged timing | None | 7.22 (9.33) | 553,743 | 39.1 | 1.000 / 1.000 |
| Submitted fixed-3-s timing† | None | 9.08 (13.00) | 537,084 | 37.9 | .995 / .963 |
| Response model | Press-to-release response boxcars | 10.48 (20.85) | 613,035 | 43.3 | .966 / .926 |
| Previous-hold model | Preceding-hold modulators | 7.36 (9.55) | 444,042 | 31.3 | .969 / .870 |

*Note.* Whole-brain extents are positive DT > TT effects after 5,000-permutation TFCE correction at *p_FWE_* < .05; *N* = 27. Spatial correlations and Dice coefficients are computed against the primary model: correlations compare unthresholded group *t*-maps within the common analysis mask, and Dice coefficients compare thresholded significant masks. VIF summaries are across participant-runs for the Time-DT canonical-HRF regressor.

Placing the cue and encoding regressors at their nominal times left the group map essentially unchanged (*r* = .995, Dice = .963; 37.9% of the analysis mask). The switching effect, therefore, does not depend on frame-accurate event timing, although the logged times did reduce collinearity among the cue-locked regressors (median Time-DT VIF 7.22 versus 9.08).

The effect was not attenuated consistently by response control. Explicit response boxcars increased the significant extent from 39.1% to 43.3% of the analysis mask, whereas the preceding-hold modulator reduced it to 31.3%; both controlled maps remained highly similar to the primary map (*r*s = .966 and .969; Figure S2). The response model had higher collinearity with the Time-DT regressor than the primary and previous-hold models, reinforcing its role as a sensitivity analysis. Across participants, the DT − TT response-latency difference was unrelated to the mean neural contrast within the primary significant mask, *r* = .055, *p* = .784; none of the four executive-control ROI associations reached significance (all *p* ≥ .181). These analyses show that the widespread DT > TT pattern is stable to the modeled response-timing variables. They do not identify the pattern as purely attentional because switch status remains inseparable from the preceding task domain in this design.

**Table S4.** Positive-control ROI tests of the previous-hold covariate on current time-cued trials.

| **ROI** | **Definition** | ***t*(26)** | ***d_z_*** | **Raw *p*** | **FDR *q*** |
| --- | --- | --- | --- | --- | --- |
| Left precentral gyrus | 8-mm sphere, MNI -38, -22, 56 | -2.44 | -0.47 | .022 | .033 |
| SMA/pre-SMA | 8-mm sphere, MNI 0, 10, 50 | -2.98 | -0.57 | .006 | .018 |
| Right putamen | 8-mm sphere, MNI 24, 4, 4 | -0.46 | -0.09 | .65 | .65 |

*Note.* Values are canonical-HRF coefficients for the previous response time from the previous-hold model; the temporal derivative received zero contrast weight. The tests are two-sided one-sample *t*-tests across *N* = 27, with Benjamini–Hochberg FDR correction across the three diagnostic ROIs. They establish that the added covariate captures motor-related variance in precentral and medial premotor cortex. The separate DT > TT map-comparison results in Table S3 show that the transition contrast remains spatially stable after adding this covariate; neither result demonstrates complete motor independence.

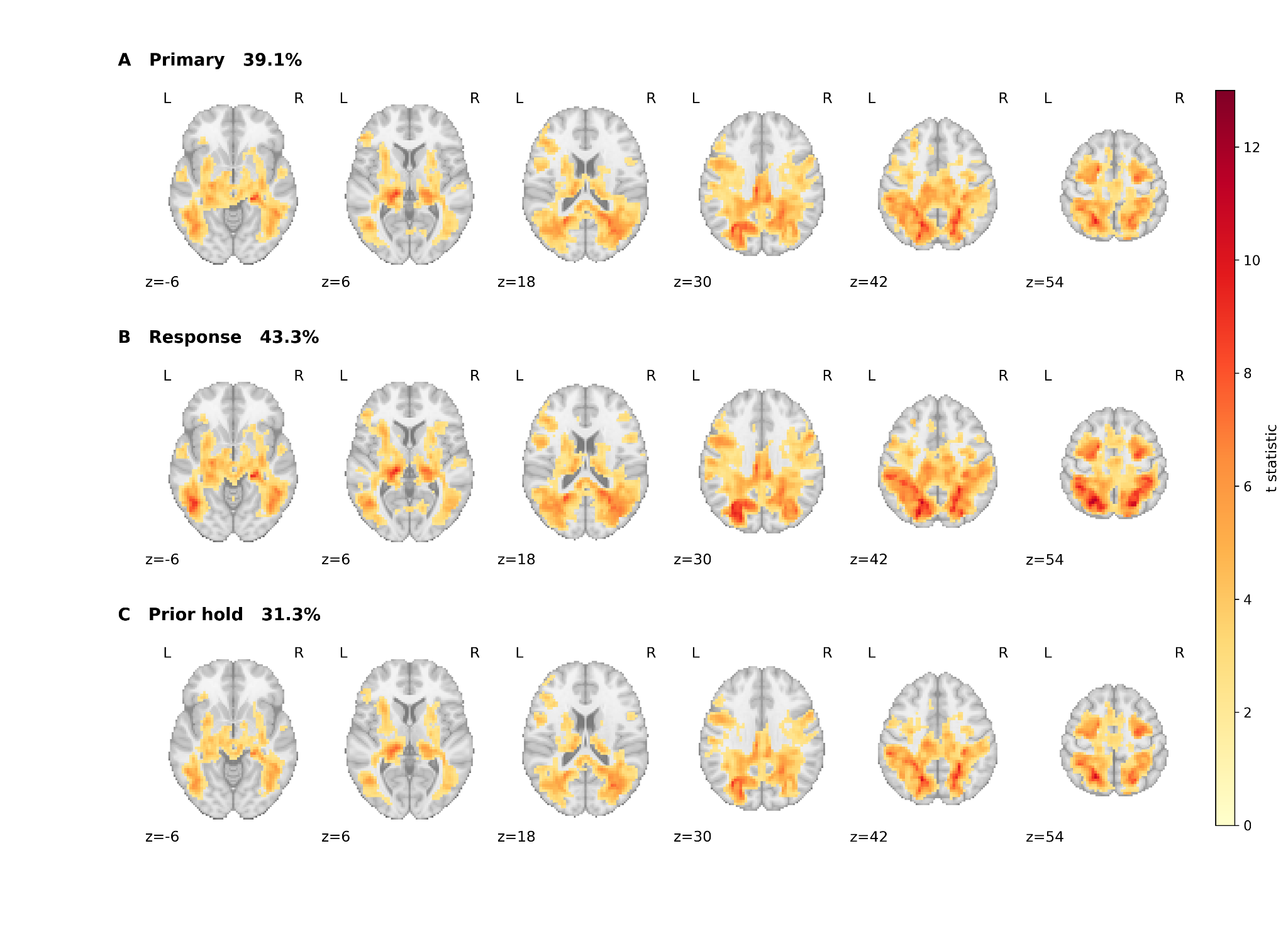

**Figure S2.** Sensitivity of the positive DT > TT effect to measured motor-carryover controls. Thresholded *t*-statistics are shown at the same axial coordinates and on a common scale after 5,000-permutation TFCE correction (*p_FWE_* < .05; *N* = 27). (**A**) Primary model with flip-logged cue timing and the trial-specific presentation-duration boxcar. (**B**) Response model with observed press-to-release boxcars. (**C**) Previous-hold model with preceding hold duration entered as a competing cue-locked modulator. Percentages denote the significant proportion of the common analysis mask.
